# Metabolite Fingerprinting for Phenotypic Screening by Infrared Matrix-Assisted Laser Desorption Electrospray Ionization Mass Spectrometry

**DOI:** 10.1101/2023.12.20.572616

**Authors:** Alena N. Joignant, Fan Pu, Shaun M. McLoughlin, James W. Sawicki, Andrew J. Radosevich, Renze Ma, Jon D. Williams, Sujatha M. Gopalakrishnan, Nathaniel L. Elsen

**Affiliations:** Discovery Research, AbbVie Inc., 1 North Waukegan Road, North Chicago, Illinois 60064, United States

**Keywords:** IR-MALDESI, phenotypic screening, drug discovery, cell-based assay, glutaminase, mass spectrometry

## Abstract

The adoption of mass spectrometry for high-throughput screening in drug discovery has become increasingly prevalent and has enabled label-free screening against diverse targets. Cellular assays for phenotypic screening, however, are primarily conducted by microscopy as there remain many challenges associated with conducting phenotypic screens via ultra-high throughput mass spectrometry.

Following a simple on-plate extraction, infrared matrix assisted laser desorption electrospray ionization (IR-MALDESI) was employed to directly sample the cell lysate at a speed of one second per sample. A549 cells were treated with compounds identified as hits in literature including a recently reported glutaminase cellular screen. Among the test compounds, there were confirmed glutaminase inhibitors, proposed nuisance compounds, and cell-active but enzyme-inactive compounds. Filtered data were further processed in R for dimensionality reduction and unsupervised clustering.

Though it was observed that all compounds affected the intracellular conversion of glutamine to glutamate, there were clear metabolic differences between the biochemically active compounds and the off-target hits. Moreover, two nuisance compounds were observed to cluster separately from the confirmed glutaminase inhibitors in the observed metabolite fingerprints.

This proof-of-concept work establishes a complete workflow that enables highthroughput mass spectrometry-based phenotypic screening. The methods proposed herein, at the throughput enabled by IR-MALDESI, could offer a new avenue for discovery of novel drugs.

## INTRODUCTION

One of the primary challenges in drug discovery is the high drug attrition rate that contributes to the high cost and effort associated with bringing life-saving therapeutics to the hands of patients.^1^ While many variables factor into this attrition, a key aspect is a lack of therapeutic targets that can be pharmacologically modulated within an acceptable safety window. Therefore, there is significant work surrounding the optimal way to identify potential new drug targets and therapeutics.

Presently, the main paradigms of early drug discovery can be distilled into two categories: target-based and phenotypic drug discovery.^2^ Target-based approaches maintain their popularity due to an increase in knowledge of the molecular drivers of many pathologies, in large part owing to revolutions in genome editing and nextgeneration sequencing. However, target-based methodologies tend to be reductionist, concentrating on singular protein interactions within *in vitro* biochemical and cellular assays. Additionally, the number of targets deemed “druggable” makes targeted approaches inefficient relative to the expanse of the human proteome, and the identification of novel targets is an unmet goal. ^3^ While these methods have contributed to bringing dozens of small molecule therapeutics to patients, reducing the early discovery process to single targets significantly limits the potential to discover new or multi-target mechanisms of action (MoAs).

Phenotypic screens, in contrast, expose compounds to a cellular disease model, and the resulting phenotypes can be used to identify potential therapeutics. This methodology provides more biologically-relevant information in a target-agnostic manner.^4^ Phenotypic screening, generally considered to be a classical method of drug discovery, has recently seen resurgence in the field due to significant technological advancements for both target identification and small-molecule discovery. Contrary to most target-based approaches, phenotypic screens offer the ability to deliver novel and first-in-class therapeutics.^5,6^ Many phenotypic assays are accomplished by fluorescence microscopy or flow cytometry, and investigate multiple broader phenotypes such as cytotoxicity and cell-cycle arrest.^2,7^ The past decade has seen the rise of cell painting via high-content screening driven by significant advancements in data analysis. In cell painting approach, several dyes are used to stain various cellular compartments. Sophisticated data analysis workflows perform automated feature extraction of hundreds of cellular attributes for clustering phenotypes by machine learning or multivariate analysis.^8^ After this clustering, the challenge remains to elucidate the targets and MoAs associated with the phenotypes, but this burden decreased with recent advancements including genome-wide CRISPR screens.^4–6^

While the observation of morphological phenotypes has advanced phenotypic screens, it is understood that the observable endpoints resulting from compound treatment are caused by changes in the underlying metabolic pathways.^9,10^ Many have explored mass spectrometry (MS)-based approaches such as metabolomics as ways to accomplish phenotypic drug discovery by monitoring metabolites and the associated activation of biological pathways corresponding to observed phenotypes.^7,10^ Beyond this, advanced data analysis methods are able to resolve MoAs based on the relevant metabolic pathways using machine learning classifiers.^11–13^ While this could be accomplished with high metabolite coverage using methods like liquid chromatography MS (LC-MS), the throughput requirements of large library screening necessitates direct infusion or direct-analysis methods.^14^ Matrix-assisted laser desorption/ionization (MALDI) MS is one such method that is commonly used for cell classification^15,16^ and has shown aptitude for cell-based phenotypic assays not limited to those investigating stress^17^, fatty acid synthase inhibition^18^, and kinase inhibition^19^.

Infrared matrix-assisted laser desorption electrospray ionization (IR-MALDESI) is a hybrid ambient ionization source that combines the advantages of electrospray ionization (ESI) and matrix-assisted laser desorption/ionization (MALDI), making it amenable to the direct analysis of diverse analytes.^20^ In IR-MALDESI, the sample is pulsed with an infrared laser, resulting in excitation and desorption of the sample. The plume of sample is intercepted by the electrospray cone, from which analyte ions are generated in an ESI-like mechanism. ^21^ While IR-MALDESI is a mature ionization source, it was only in the last five years that it has cemented its place in high-throughput screening for early drug discovery. ^22,23^ The speed of IR-MALDESI high-throughput screening is generally limited by the resolving power of the instrument, up to speeds of 22 samples per second, at which point there is less than 1% sample carryover.^24^ MALDESI has therefore become instrumental in target-based biochemical assays and screens due to its efficiency and sensitivity.

A workflow was recently developed to accomplish high-throughput cellular pharmacodynamics of glutaminase inhibitors using IR-MALDESI by directly analyzing cell lysate.^25^ As glutaminase governs the conversion of glutamine to glutamate, the relative abundances of these two metabolites were monitored to determine the activity of glutaminase inhibitors relative to control. The study culminated in a primary screen of more than 100,000 compounds. Aside from glutamine and glutamate, this method also resulted in the putative identification of approximately 400 other metabolites. Therefore, it was postulated that the method could be adapted to accomplish high throughput metabolite fingerprinting and MS-based phenotypic screening. Herein, a similar cell culture and metabolite extraction protocol was adopted to perform metabolite fingerprinting of treated cell culture to enable phenotypic screening by IR-MALDESI-MS.

## RESULTS AND DISCUSSION

### Development and Optimization of Metabolite Extraction

A schematic of the metabolite extraction and data analysis workflows is shown in **Figure 1**. It was observed that cell overcrowding and ion suppression increased when plating more than 5,000 cells, since an increase in cells generally results in a broad increase in metabolite abundances that may suppress lower-abundance species. Therefore, 5,000 cells per well became the optimal cell density for this method (**Figure S1**). After incubation with compound, the cell media was removed by washing twice with 150 mM ammonium acetate, which is MS-compatible. Washed cells were extracted in 50 µL of 20% methanol in water containing the two internal standards, a procedure that was previously found to work in this format and serves as the IR-MALDESI solvent for analysis. To prevent uneven evaporation across the plate, the plates were covered with silicone lids and left to extract for thirty minutes at 4°C. It was observed that higher ion signal and lower variability was achieved when 20 µL of the cell lysate was transferred to Proxi plates prior to IR-MALDESI-MS analysis (**Figure S2**).

**Figure 1.**
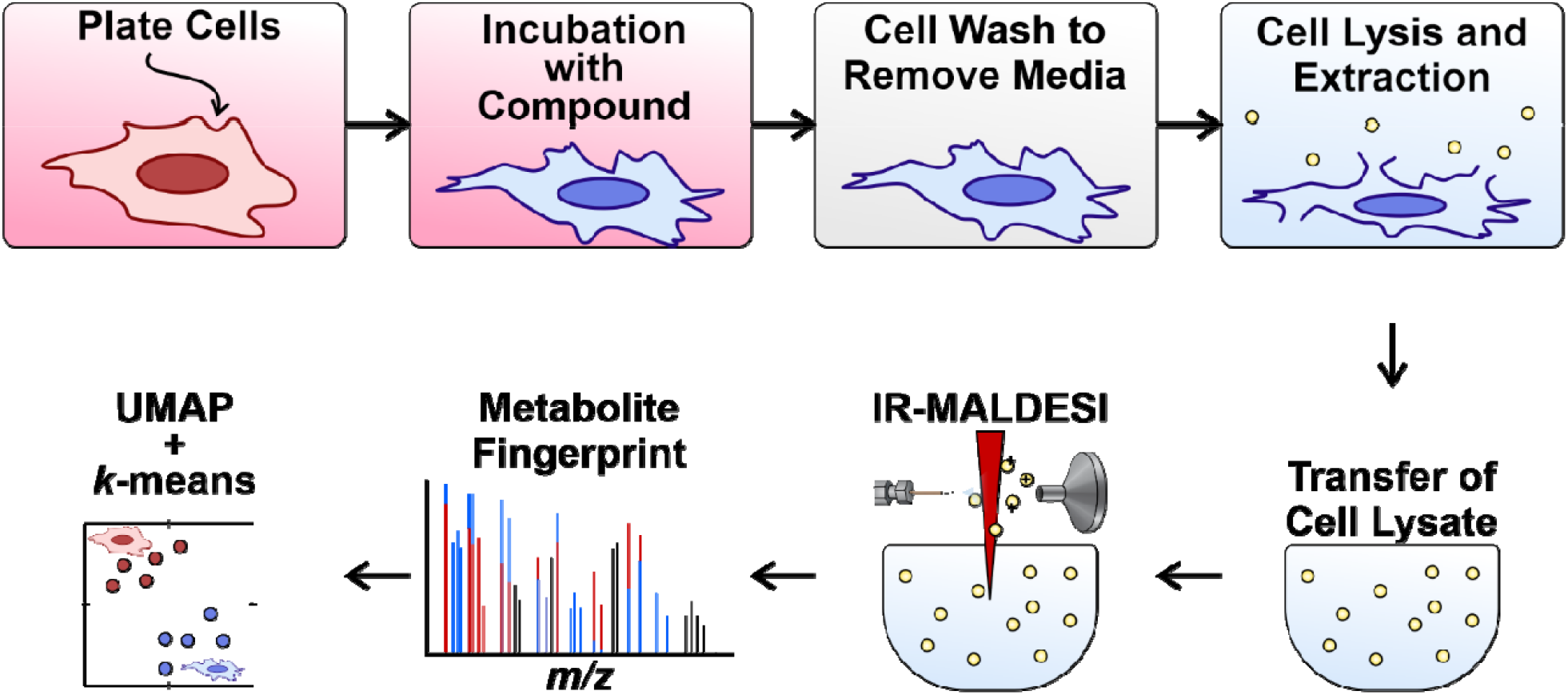
Outline of the experimental workflow optimized for metabolite fingerprinting.

### Successful Clustering of Glutaminase Inhibitors from Control

Preliminary experiments consisted of preparing two plates of cells and treating half of each plate with a confirmed glutaminase inhibitor. The cells were left to incubate and adhere overnight (16 hours). The following day, the cell media was evacuated using a Blue Washer “Lightspin” protocol (BlueCatBio). Evacuated cells were inspected under a light microscope to ensure that little to no cell loss was observed throughout the protocol. Half of each plate was treated with 20 µL of 10 µM of the glutaminase inhibitors CB839 or BPTES in media. An equal volume of control media was added to the other half of each plate. The cells were incubated with compound for four hours prior to metabolite extraction and IR-MALDESI-MS analysis in positive ion mode. **Figure S3** illustrates the efficacy of both compounds after this four-hour incubation. Abundance matrices were prepared and input into R for Uniform Manifold Approximation and Projection (UMAP), which was recently developed by McInnes *et al*. and described elsewhere.^26^ The UMAP plots for half-plated CB839 and half-plated BPTES are shown in **Figure 2** and **Figure 3**, respectively. *K*-means clustering was applied on the raw UMAP output to assign each well (point on the UMAP) to a cluster as a way of evaluating how effective and accurate the clustering was for each group in an unsupervised manner. Confusion matrices are included for each UMAP, showing how many wells were classified into which cluster. This method is generally used for supervised techniques to determine accuracy, but since the treatment in each well was known, it is used in this case to evaluate the accuracy of the unsupervised clustering method. The CB839-treated wells were able to separate from control with 94% accuracy. A small cluster of control wells can be seen in the bottom right corner of the UMAP and are an artifact of signal dropout in the control half of the plate. The BPTES wells were able to separate from control with 97% accuracy, as determined by the confusion matrix.

**Figure 2.**
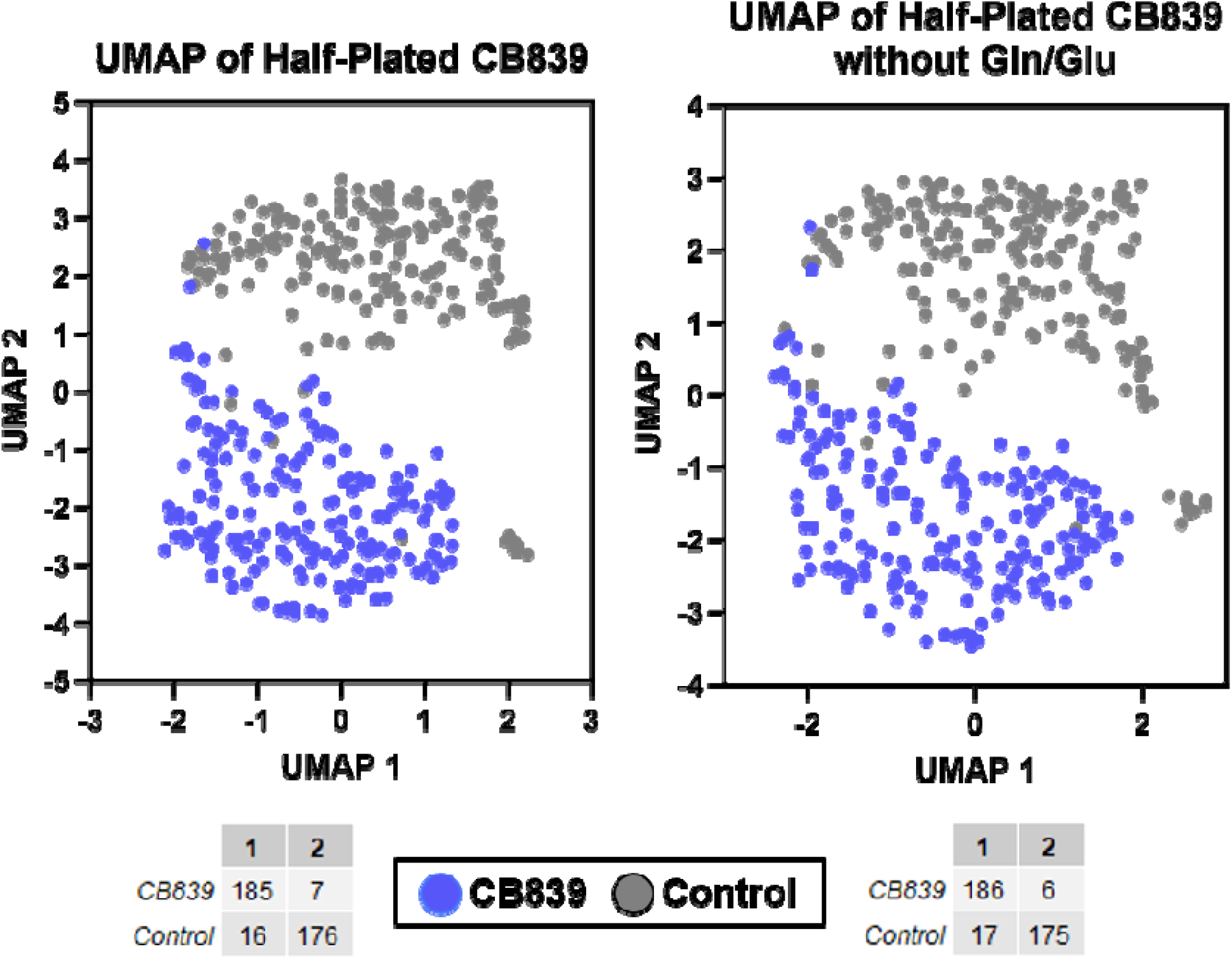
True clustering of half-plated CB839 with (left) and without (right) annotations corresponding to glutamine or glutamate. UMAP was conducted with default minimum distance and 30 nearest neighbors. *k*-means clustering was able to delineate the two groups with 94% accuracy in both trials based on the confusion matrices.

**Figure 3.**
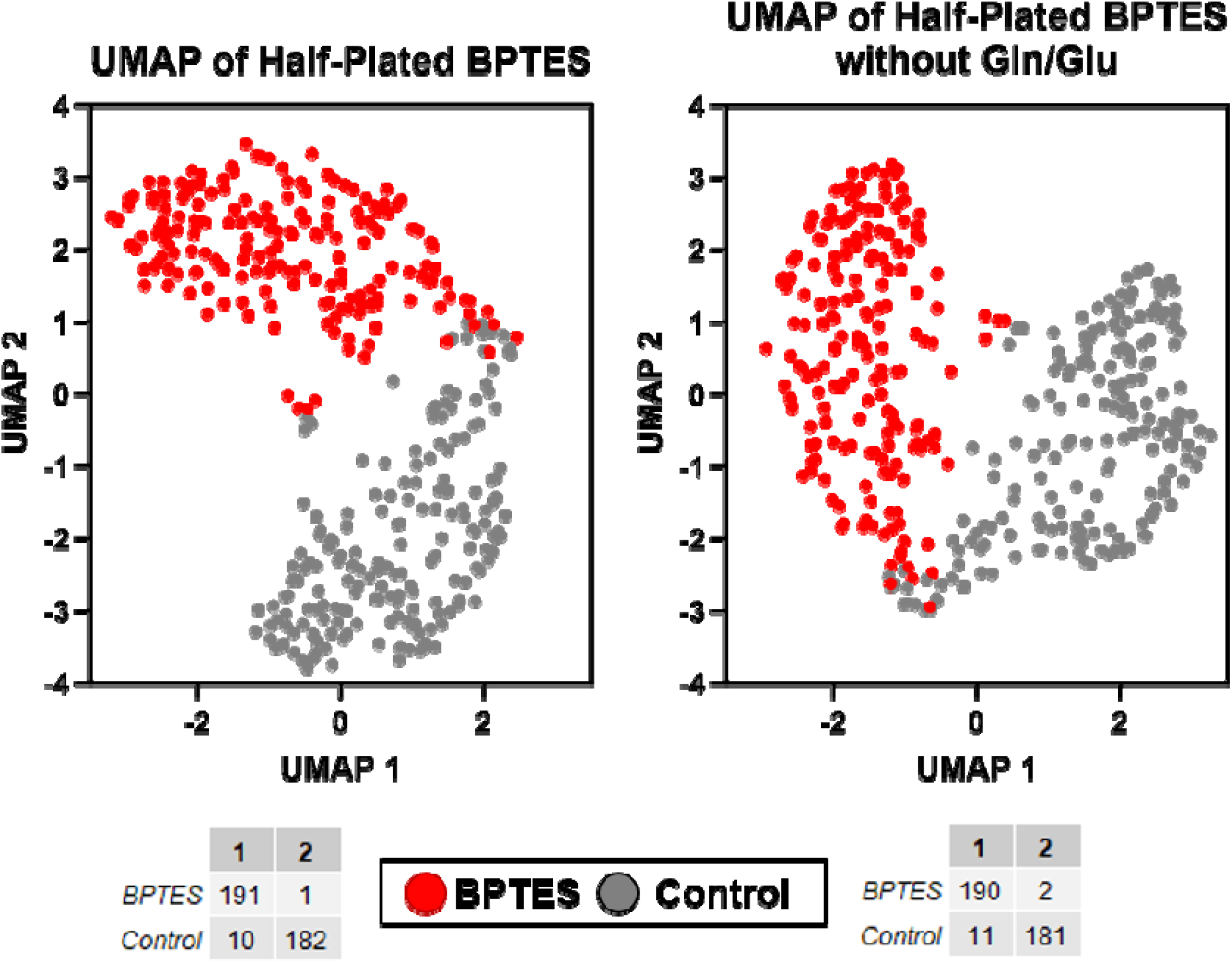
True clustering of half-plated BPTES with (left) and without (right) annotations corresponding to glutamine or glutamate. UMAP was conducted with default minimum distance and 30 nearest neighbors. *k*-means clustering was able to delineate the two groups with 97% (left) and 94% accuracy (right).

The annotations that correspond to glutamine or glutamate were removed from the dataset, and the data was re-analyzed to ensure that prior clustering was not primarily due to the relative abundances of glutamine and glutamate. The clustering accuracy in the CB839 data set remained 94%, indicating no loss of accuracy. The 6% of wells that were inaccurately clustered likely suffered from signal variance, which would negatively affect relative abundance of metabolites and therefore clustering. There was only a small decrease in accuracy upon removal of glutamine and glutamate from the BPTES data set, but this accuracy remained high at 94%. This suggests that other metabolites must be contributing to the separation of treatment from control, indicating that this method incorporates more of the “metabolite fingerprint”, as in metabolites other than those directly impacted by enzyme modulation.

### Clustering Active, Nuisance, and Inactive Glutaminase Compounds

After confirming the ability to cluster half-plated cell treatments, the method was further tested for its ability to distinguish compound MOAs using a total of six compounds. The list includes the confirmed glutaminase inhibitors CB839 (EC_50,cellular_ = 0.004 µM) and BPTES in addition to four compounds previously identified as cellular hits in a glutaminase pharmacodynamics assay.^25^ Two of these compounds were active in both the cellular screen and the biochemical assay but were found to not bind to glutaminase by surface plasmon resonance (data not shown), and were later found to be nuisance compounds (Nuisance 1 and Nuisance 2, EC_50,cellular_ = 9.659 and 1.568 µM, respectively). To evaluate the possible nuisance mechanism, 100 µM of each six compounds were incubated with 0.5 mM Tris(2-carboxyethyl)phosphine (TCEP) for 24 hours, then oxidation of TCEP was analyzed using IR-MALDESI-MS. Such MS based assay has been reported for detection of redox cycling compounds.^27^ The results are shown in **Figure S4**. The two nuisance compounds showed moderate levels of redox activity, which likely caused non-specific inhibition of the enzyme. The final two compounds included in this experiment were previously observed to be active in the cellular screen but not in the biochemical assay and are considered off-target (Inactive 1 and Inactive 2, EC_50,cellular_ = 2.321 and 1.405 µM, respectively). To further confirm intracellular target engagement, or lack thereof, for the 6 compounds profiled, a cellular thermal shift assay (CETSA) was performed. Gratifyingly, the two confirmed glutaminase inhibitors, CB839 and BPTES, showed a marked stabilization of GLS1 at 46°C while the other four compounds showed no stabilization (**Figure S5)**.

An Echo 555 (Beckman Coulter) was used to dispense compounds into empty cell culture plates prior to plating cells. In this case, 2.5 or 25 nL of 10 mM compound in DMSO were dispensed so that the final compound concentration was 1 or 10 µM in 25 µL of media. The 24-hour incubation began immediately upon cell plating. A replicate plate was incubated in parallel to assess cell viability to account for metabolite abundances modulated only due to relative viability. It was observed that the compounds had minor differences in their impact on cell viability at 10 µM, indicating that the cell counts in each well were comparable and did not influence overall metabolite abundances (**Figure S6**). The efficacy of all six compounds at this dose and incubation time was demonstrated by plotting the average normalized ratio of glutamine to glutamate in the mass spectra (**Figure S7**). The compounds that had the largest effect on the metabolite ratio were CB839, BPTES, and the two nuisance compounds. In contrast, the two enzyme-inactive compounds produced only slightly altered metabolite ratios relative to the untreated cells.

Figure 4. displays the UMAP of all the compounds in both positive and negative ion mode. The UMAP plots show a clear separation of enzyme-active treatments from inactive treatments and control. The smaller groupings, of <10 wells, that are separate from the main clusters correspond to variance in ion abundance affecting those wells (*i*.*e*., hotspots or signal decreases). Since the treatments are generally differentiated by their relative abundances of each metabolite, variance in ion abundance is a large obstacle for unsupervised compound clustering. The enzyme-inactive hits were much less effective than true inhibitors at modulating metabolite ratios, so it is unsurprising that these compounds did not separate from control in projections that are based on metabolite abundances. This demonstrates the promise of this method to resolve enzyme-active true hits from enzyme-inactive false hits in future screens.

**Figure 4.**
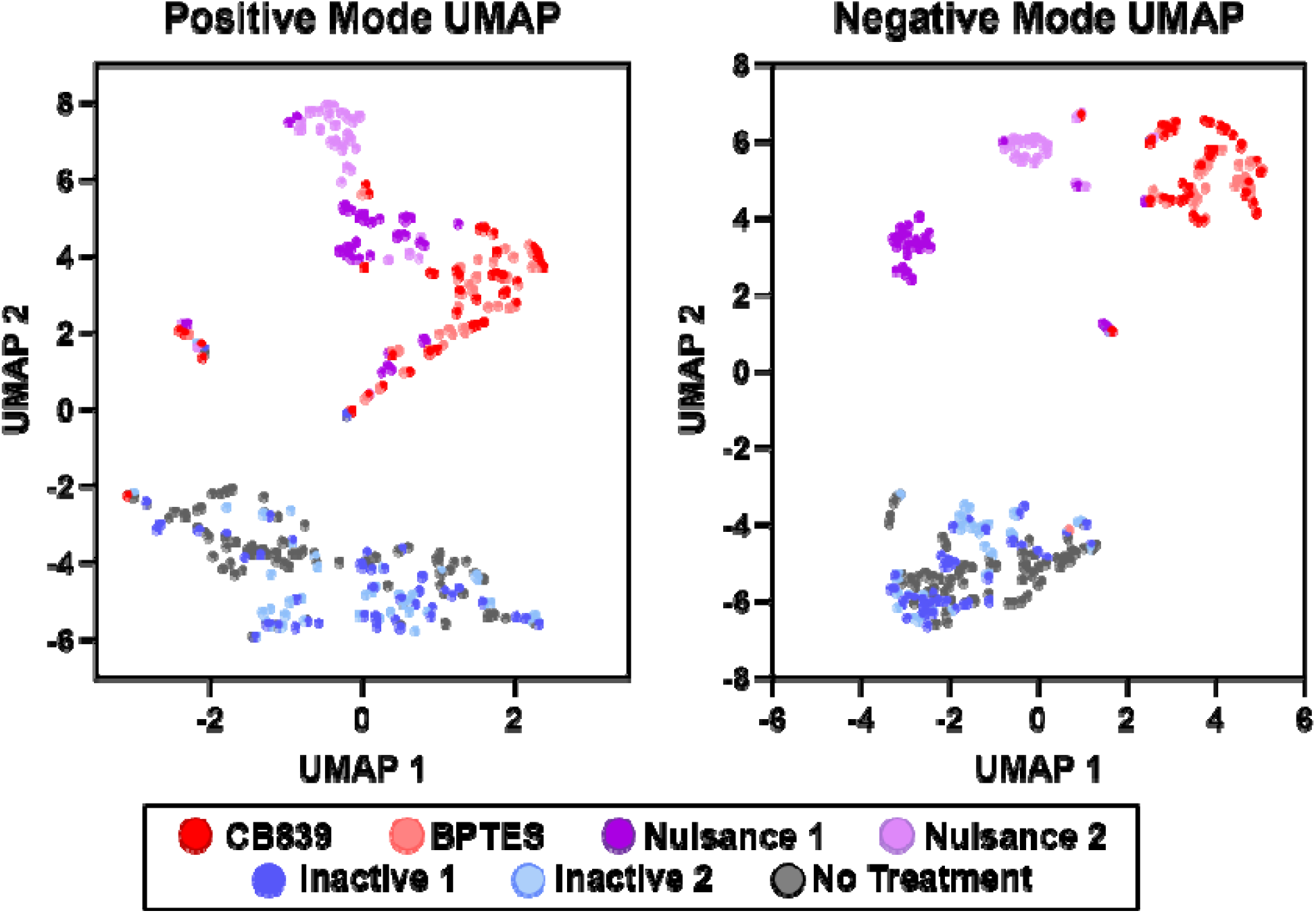
UMAP of all glutaminase compounds and control in positive mode (left) and negative mode (right). Wells are colored by treatment (true values). UMAP was conducted with a minimum distance of 0.001 and 5 nearest neighbors. Due to the irregular shape of the UMAP clusters in each case, k-means was ineffective at evaluating the accuracy of separation.

The two confirmed glutaminase inhibitors, CB839 and BPTES, completely overlap in the UMAP plot. This indicates that the metabolite fingerprints for these compounds are indistinguishable. The nuisance compounds show some separation from the confirmed inhibitors as well as from each other, suggesting metabolic differences that do not exist between CB839 and BPTES. This provides additional evidence that may confirm the off-target nature of these nuisance compounds, since different metabolites are impacted due to these treatments.

It was observed that the compounds are better separated in negative mode, which is likely because most ions that contribute to the metabolic differences between treatments were identified primarily in negative mode. Some notable metabolites that were identified as contributing to this separation are summarized in **Figure 5**. Glutaminase inhibition is a known modulator of cell metabolism through interference with downstream TCA cycle intermediates alpha-ketoglutarate, malate, fumarate, and citrate. Additionally, the conversion of glutamine to glutamate makes nitrogen available for downstream amino acid synthesis.^28^ The decrease in glutathione abundance is also expected due to the decrease of the glutamate precursor. Aspartate, which is closely linked to glutamate, was also shown to decrease after glutaminase inhibition. ^29^ The two nuisance compounds show differing abundance of citric acid relative to the confirmed inhibitors, indicating that they may also be impacting the TCA cycle in an alternative manner. Other ions that differentiate the two nuisance compounds 1 and 2 are threonic acid and gluconic acid. Interestingly, treatment with the two nuisance compounds resulted in slightly higher abundance of arginine in the cells, while the other compounds had comparable abundance to control. In most cases, the enzyme-inactive compounds produce comparable metabolite abundances to the untreated cells, further explaining their inability to separate from control in the UMAP.

**Figure 5.**
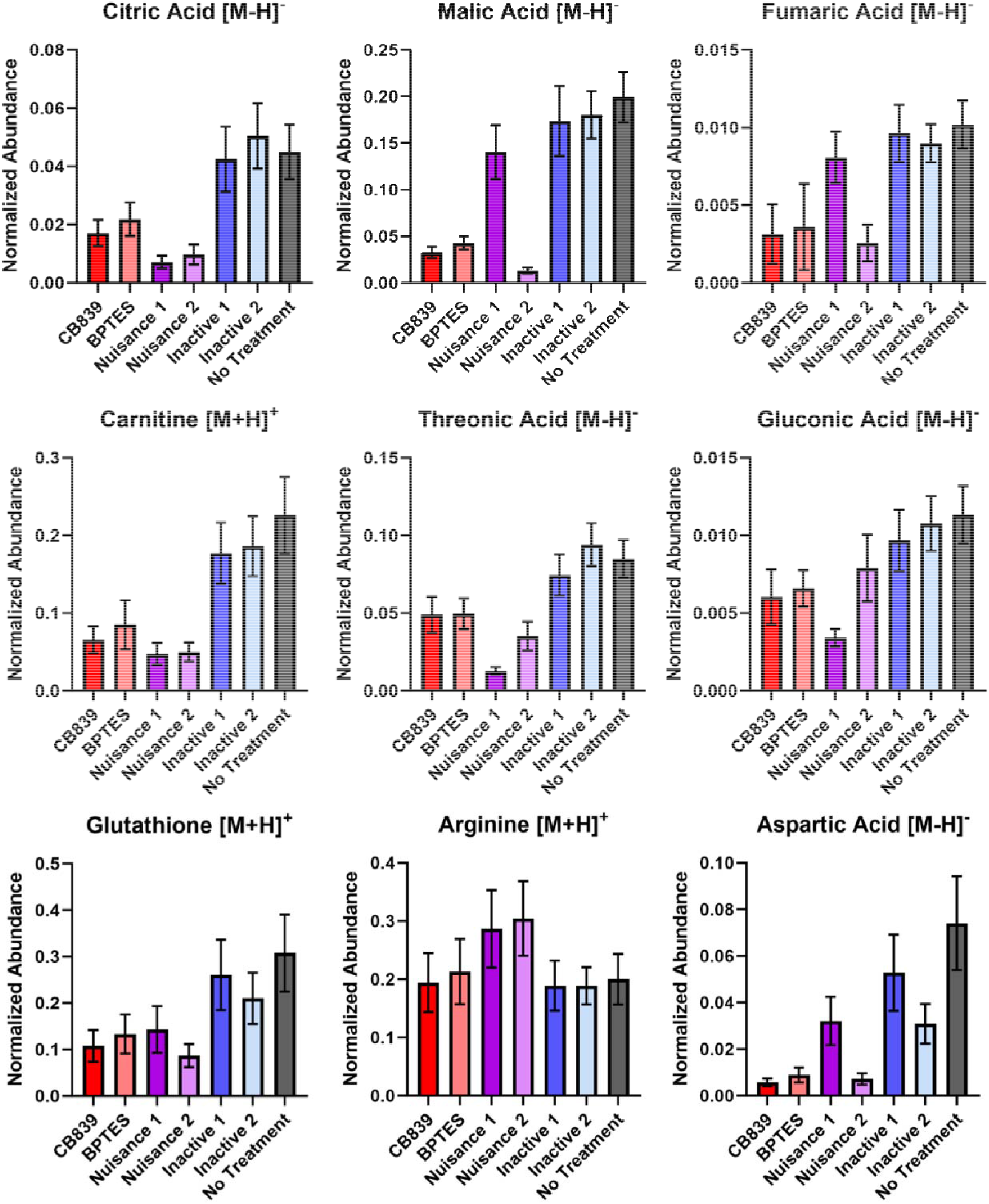
Normalized average abundances of metabolites that contribute to differences between the compounds: citric acid (*m/z*: 191.0197, [M-H]^-^), malic acid (*m/z*: 133.0143, [M-H]^-^), fumaric acid (*m/z*: 115.0037, [M-H]^-^), carnitine (*m/z*: 162.1125, [M+H]^+^), threonic acid (*m/z*: 135.0299, [M-H]^-^), gluconic acid (*m/z*: 195.051, [M-H]^-^), glutathione (*m/z*: 308.0911, [M+H]^+^), arginine (*m/z*: 175.1189, [M+H]^+^), and aspartic acid (*m/z*: 132.0302, [M-H]^-^). Positive ions were normalized to ^13^C_3_-caffeine, and negative ions were normalized to succinic acid-d_4_. Errors bars correspond to standard deviation.

The methods described herein work to unbiasedly cluster compounds without the need to pre-train supervised machine learning models. Therefore, this work may be expanded to investigate diverse compounds beyond those impacting glutaminase. Additionally, it was demonstrated that compounds can not only be clustered based on phenotype, but that differing MoAs may be resolved. These differences were not identified in the previous screen monitoring only glutamine and glutamate, illustrating the importance of the “metabolite fingerprint” in elucidating the mechanisms of compounds that achieve similar phenotypes.

Future studies investigating many more compounds would likely require clustering of multiple plates of data. In this case, the impact of batch effects on the ability to cluster compounds must be identified. Additionally, the mitigation of signal variability is crucial to maintain clustering accuracy when adding more diverse compound treatments. To advance the data analysis methods, it is possible to incorporate supervised machine learning and conduct recursive feature elimination to identify metabolites that are the most predictive of a certain phenotype. To accomplish this, training data must be collected for known MoAs. If this was done, it may elucidate biomarkers for the desired phenotypes that could be used for a larger primary screen at the expense of creating a more specialized model.^7,11^ Finally, the long-term success of MS-based phenotypic screening would be further enabled by integrating multi-omics to survey the cellular phenotypes with a systems biology approach and better-resolve the MoAs and potential targets of cell-active compounds simultaneously.

## CONCLUSION

This work enables the future development and application of IR-MALDESI-MS phenotypic screening to improve our understanding of the behavior of compounds in biological context. The use of metabolites in this work was effective in distinguishing compounds by their metabolite fingerprint. Specifically, these methods indicate promise in separating active compounds from inactive compounds, as well as distinguishing off-target or nuisance compounds from confirmed hits. Using UMAP and *k*-means clustering allows this method to be applied to diverse mechanisms of action without the need to train supervised machine learning models prior to application. Future assay and data analysis development will likely see this work lead to a routinely used tool in early drug discovery.

## METHODS

### Cell Culture

A549 cells (ATCC) were used throughout this work. The cells were cultured in 150 mm^3^ flasks (Corning) in DMEM media + 10% fetal bovine serum (Sigma Aldrich) and 1% antibiotic-antimycotic (Gibco). The same cell lineage was used in all experiments with undetermined passage number. The cell culture was split upon reaching 80% confluence. To prepare experimental plates of cells, the cell monolayer was detached using 0.25% Trypsin-EDTA (Gibco), and the cells were counted using an i-cell XR cell viability analyzer (Beckman Coulter).

For the CETSA assay, after the cells were detached with 0.25% trypsin, the cells were washed with HBSS buffer (Gibco) and resuspended to 20 × 10^6^ cells/mL with HEPES buffer (pH 7.4) containing 138 mM NaCl, 5 mM KCl, 2 mM CaCl2, and 1 mM MgCl2.

For other assays, the cell solution was diluted using fresh media, and 5,000 cells per well were plated on cell culture plates (Greiner bio one) using a Multidrop Combi dispenser (Thermo Fisher Scientific).

### Metabolite Extraction

After incubation with compound, cell plates were washed twice with 150 mM ammonium acetate on a Blue Washer (BlueCatBio). Washed cells were extracted with 50 µL extraction solvent composed of 20% MeOH in water and including stable isotope-labelled caffeine (^13^C_3_, 8 µM, Cambridge Isotope Laboratory) and deuterated succinic acid (d_4_, 1 µM, Sigma Aldrich) as internal standards in positive and negative mode, respectively. Immediately following the dispensing of the extraction solvent, the cells were shaken by the Combi for 10 seconds to induce cell lysis. Then, the Greiner plates were capped with silicone plate lids (Axygen) and left to extract at 4°C in the fridge. After 30 minutes, 20 µL of the cell lysate in each well were transferred to a Proxi plate by a Beckman NX (Beckman Coulter). The plates were then promptly de-ionized (to prevent ESI spray instability) and analyzed by IR-MALDESI-MS.

### Cell Viability Assays

An extra plate was treated with compound and reserved for CellTiter-Glo cell viability assay. After the incubation with compound, CellTiter-Glo reagent (25 µL, Promega) was added to each well. The luminescence of each well was measured using the ViewLux (PerkinElmer) and were corrected by subtracting the average luminescence of wells with no cells plated.

### TCEP Assay

0.5 M TCEP (Bond-Breaker™ TCEP Solution, Thermo Fisher Scientific) was diluted to 0.5 mM in 20 mM Tris hydrochloride buffer solution (pH = 7.5, Sigma Aldrich). Test compounds were solvated in DMSO at 10 mM and dispensed into Proxi plate at 150 nL per well using Echo. 0.5 mM buffered TCEP solution was then dispensed onto the compound plate at 15 µL per well, resulting a 100 µM compound concentration. After 24 hours incubation, the plate was analyzed with IR-MALDESI-MS in positive ion mode, a mass range of *m/z* 200-400 was used. Percent oxidation was calculated by normalizing percent conversion to blanks. Percent conversion was calculated by dividing abundances of oxidized TCEP (*m/z* 267.06) by total abundances of TCEP (*m/z* 251.07) and oxidized TCEP.

### IR-MALDESI

The details of the IR-MALDESI-HTS platform have been described previously.^20,24^ A 2970 nm wavelength solid-state laser (JGMA Inc., Burlington, MA) was fired into the sample well in three laser bursts of two pulses each. The electrospray solvent consisted of 80% methanol in water with an additional 0.1% formic acid and 100 nM unlabeled caffeine. Mass spectra for phenotypic screening were acquired on an Orbitrap HFX (Thermo Fisher Scientific, Bremen, Germany) with a resolving power setting of 120,000 FWHM and a 20-ms injection time. A mass range of *m/z* 100-400 was used in all experiments. MS1 spectra were acquired by combining three microscans, each microscan corresponding to one laser sampling event in the well. The throughput of the method was determined by the three sampling events per well, resulting in an approximate 6 minutes per plate.

## Data Analysis

Raw data files from MALDESI were directly uploaded to MSiReader v2.51 BioPharma with no file conversion (MSI Software Solutions, LLC; Raleigh, NC). Spectra from untreated A549 cell lysate were annotated using the RefMet database in Metabolomics Workbench with a mass tolerance of 5 ppm (https://www.metabolomicsworkbench.org/). Annotations corresponding to background ambient ions were removed. Only the annotation with the smallest mass error for each experimental peak was included in the final annotation list in each mode. The annotations lists used in this work are included as a supplemental xlsx file. Isomers were not resolved. The MSiExport tool in MSiReader was used to export the normalized abundance of the annotations in each well as an Excel spreadsheet.

Dimensionality reduction was done using Uniform Manifold Approximation and Projection (UMAP) in R v4.2.3 and R Studio (R Core Team 2021). The abundance matrix output from MSiExport was reformatted as a data frame for input into R to contain an additional column for compound ID in each well. The normalized well abundances for each annotation were scaled and centered using the “scale()” function in R. If not otherwise specified, default “umap()” parameters were used (https://github.com/tkonopka/umap). Clustering was done on the “umap$layout” data frame using k-means clustering in R with an “nstart” of 20 and otherwise default parameters. A .csv file was generated that includes the UMAP coordinates, well ID, compound ID, and cluster number. An exemplar code used in this project is publicly available at github.com/anjoigna/IR-MALDESI-MS-PS.

### CETSA assay

Resuspended A549 cells were incubated with either compounds or DMSO vehicle for 1 hr at 37 °C. The final concentration of the compounds was 10 _μ_M. Following the incubation, an aliquot of 20 _μ_L of each of the cell samples were taken out to be heated for 3 min at one of 6 temperatures that were equally spaced at 3 °C intervals between 40 and 55 °C. Once the cells cooled to room temperature, they were lysed for by 3 cycles of liquid nitrogen freeze-thaw with 1X protease and phosphatase inhibitor cocktail (Thermo Fisher) added. After the lysis, the samples were centrifuged at 30,000 × g for 20 minutes at 4 °C. The supernatant containing soluble proteins were collected after the centrifugation and were subjected to immunoblotting analysis using WesTM (Bio-Techne) simple western instrument.

## Supporting information

A549 Metabolomics Workbench Annotations

## SUPPORTING INFORMATION

- Cell density titration (Figure S1), extraction transfer improvements (Figure S2), half-plated data compound efficacy (Figure S3), TCEP assay results (Figure S4), CETSA results (Figure S5), cell viability results (Figure S6), and compound efficacy for all compounds (Figures S7)
- Putative annotations and metabolite list used for clustering (xlsx)

## ACKNOWLEDGEMENTS

The authors thank the following AbbVie employees: Sanjay Panchal, Brett Bruckner, Chris Ling, and Scott Warder for their assistance in realizing this work. The authors also thank David Muddiman (North Carolina State University) for facilitating discussion with MSI Software Solutions, LLC, concerning valuable improvements to MSiReader Pro BioPharma. No Funding to disclose.

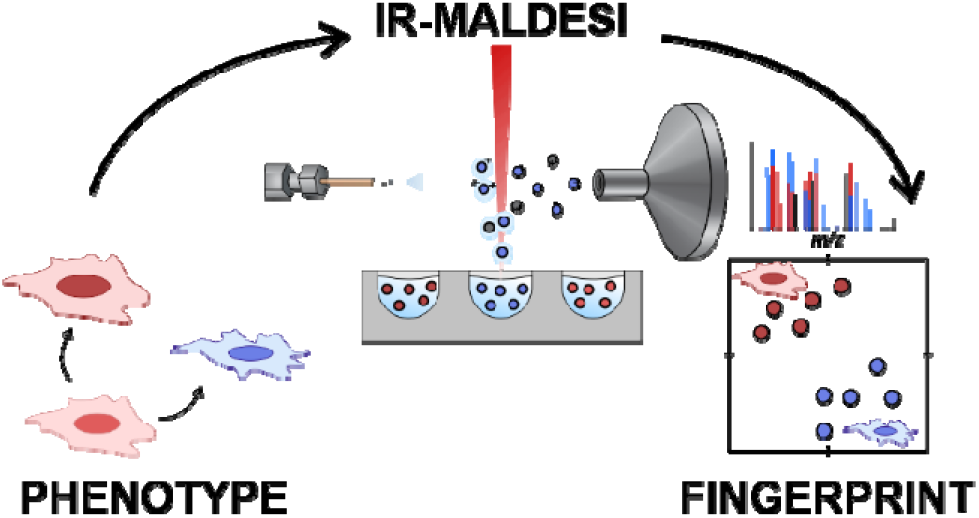

## REFERENCES

(1) Moffat, J. G.; Vincent, F.; Lee, J. A.; Eder, J.; Prunotto, M. Opportunities and Challenges in Phenotypic Drug Discovery: An Industry Perspective. Nat Rev Drug Discov 2017, 16 (8), 531–543. 10.1038/nrd.2017.111.

(2) Moffat, J. G.; Rudolph, J.; Bailey, D. Phenotypic Screening in Cancer Drug Discovery — Past, Present and Future. Nat Rev Drug Discov 2014, 13 (8), 588–602. 10.1038/nrd4366.

(3) Zheng, W.; Thorne, N.; McKew, J. C. Phenotypic Screens as a Renewed Approach for Drug Discovery. Drug Discovery Today 2013, 18 (21–22), 1067–1073. 10.1016/j.drudis.2013.07.001.

(4) Palmer, M. Phenotypic Screening. In Small Molecule Medicinal Chemistry; Czechtizky, W., Hamley, P., Eds.; Wiley, 2015; pp 281–304. 10.1002/9781118771723.ch10.

(5) Wagner, B. K.; Schreiber, S. L. The Power of Sophisticated Phenotypic Screening and Modern Mechanism-of-Action Methods. Cell Chemical Biology 2016, 23 (1), 3–9. 10.1016/j.chembiol.2015.11.008.

(6) Vincent, F.; Nueda, A.; Lee, J.; Schenone, M.; Prunotto, M.; Mercola, M. Phenotypic Drug Discovery: Recent Successes, Lessons Learned and New Directions. Nat Rev Drug Discov 2022, 21 (12), 899–914. 10.1038/s41573-022-00472-w.

(7) Dueñas, M. E.; Peltier-Heap, R. E.; Leveridge, M.; Annan, R. S.; Büttner, F. H.; Trost, M. Advances in High-throughput Mass Spectrometry in Drug Discovery. EMBO Mol Med 2023, 15 (1). 10.15252/emmm.202114850.

(8) Bray, M.-A.; Singh, S.; Han, H.; Davis, C. T.; Borgeson, B.; Hartland, C.; Kost-Alimova, M.; Gustafsdottir, S. M.; Gibson, C. C.; Carpenter, A. E. Cell Painting, a High-Content Image-Based Assay for Morphological Profiling Using Multiplexed Fluorescent Dyes. Nat Protoc 2016, 11 (9), 1757–1774. 10.1038/nprot.2016.105.

(9) Kell, D. B. Systems Biology, Metabolic Modelling and Metabolomics in Drug Discovery and Development. Drug Discovery Today 2006, 11 (23–24), 1085–1092. 10.1016/j.drudis.2006.10.004.

(10) Guijas, C.; Montenegro-Burke, J. R.; Warth, B.; Spilker, M. E.; Siuzdak, G. Metabolomics Activity Screening for Identifying Metabolites That Modulate Phenotype. Nat Biotechnol 2018, 36 (4), 316–320. 10.1038/nbt.4101.

(11) Van Oosten, L. N.; Klein, C. D. Machine Learning in Mass Spectrometry: A MALDI-TOF MS Approach to Phenotypic Antibacterial Screening. J. Med. Chem. 2020, 63 (16), 8849–8856. 10.1021/acs.jmedchem.0c00040.

(12) Zampieri, M.; Szappanos, B.; Buchieri, M. V.; Trauner, A.; Piazza, I.; Picotti, P.; Gagneux, S.; Borrell, S.; Gicquel, B.; Lelievre, J.; Papp, B.; Sauer, U. High-Throughput Metabolomic Analysis Predicts Mode of Action of Uncharacterized Antimicrobial Compounds. Sci. Transl. Med. 2018, 10 (429), eaal3973. 10.1126/scitranslmed.aal3973.

(13) Smith, M. J.; Ivanov, D. P.; Weber, R. J. M.; Wingfield, J.; Viant, M. R. Acoustic Mist Ionization Mass Spectrometry for Ultrahigh-Throughput Metabolomics Screening. Anal. Chem. 2021, 93 (26), 9258–9266. 10.1021/acs.analchem.1c01616.

(14) Zampieri, M.; Sekar, K.; Zamboni, N.; Sauer, U. Frontiers of High-Throughput Metabolomics. Current Opinion in Chemical Biology 2017, 36, 15–23. 10.1016/j.cbpa.2016.12.006.

(15) Claydon, M. A.; Davey, S. N.; Edwards-Jones, V.; Gordon, D. B. The Rapid Identification of Intact Microorganisms Using Mass Spectrometry. Nat Biotechnol 1996, 14 (11), 1584–1586. 10.1038/nbt1196-1584.

(16) Serafim, V.; Shah, A.; Puiu, M.; Andreescu, N.; Coricovac, D.; Nosyrev, A. E.; Spandidos, D. A.; Tsatsakis, A. M.; Dehelean, C.; Pinzaru, I. Classification of Cancer Cell Lines Using Matrix-Assisted Laser Desorption/Ionization Time-of-Flight Mass Spectrometry and Statistical Analysis. International Journal of Molecular Medicine 2017, 40 (4), 1096–1104. 10.3892/ijmm.2017.3083.

(17) Schwamb, S.; Munteanu, B.; Meyer, B.; Hopf, C.; Hafner, M.; Wiedemann, P. Monitoring CHO Cell Cultures: Cell Stress and Early Apoptosis Assessment by Mass Spectrometry. Journal of Biotechnology 2013, 168 (4), 452–461. 10.1016/j.jbiotec.2013.10.014.

(18) RamalloGuevara, C.; Paulssen, D.; Popova, A. A.; Hopf, C.; Levkin, P. A. Fast Nanoliter-Scale Cell Assays Using Droplet Microarray–Mass Spectrometry Imaging. Advanced Biology 2021, 5 (3), 2000279. 10.1002/adbi.202000279.

(19) Weigt, D.; Sammour, D. A.; Ulrich, T.; Munteanu, B.; Hopf, C. Automated Analysis of Lipid Drug-Response Markers by Combined Fast and High-Resolution Whole Cell MALDI Mass Spectrometry Biotyping. Sci Rep 2018, 8 (1), 11260. 10.1038/s41598-018-29677-z.

(20) Caleb Bagley, M.; Garrard, K. P.; Muddiman, D. C. The Development and Application of Matrix Assisted Laser Desorption Electrospray Ionization: The Teenage Years. Mass Spectrometry Reviews 2023, 42 (1), 35–66. 10.1002/mas.21696.

(21) Sampson, J. S.; Hawkridge, A. M.; Muddiman, D. C. Generation and Detection of Multiply-Charged Peptides and Proteins by Matrix-Assisted Laser Desorption Electrospray Ionization (MALDESI) Fourier Transform Ion Cyclotron Resonance Mass Spectrometry. J. Am. Soc. Mass Spectrom. 2006, 17 (12), 1712–1716. 10.1016/j.jasms.2006.08.003.

(22) Pu, F.; Radosevich, A. J.; Sawicki, J. W.; Chang-Yen, D.; Talaty, N. N.; Gopalakrishnan, S. M.; Williams, J. D.; Elsen, N. L. High-Throughput Label-Free Biochemical Assays Using Infrared Matrix-Assisted Desorption Electrospray Ionization Mass Spectrometry. Anal. Chem. 2021, 93 (17), 6792–6800. 10.1021/acs.analchem.1c00737.

(23) Knizner, K. T.; Guymon, J. P.; Garrard, K. P.; Bouvrée, G.; Manni, J.; Hauschild, J.-P.; Strupat, K.; Fort, K. L.; Earley, L.; Wouters, E. R.; Pu, F.; Radosevich, A. J.; Elsen, N. L.; Williams, J. D.; Pankow, M. R.; Muddiman, D. C. Next-Generation Infrared Matrix-Assisted Laser Desorption Electrospray Ionization Source for Mass Spectrometry Imaging and High-Throughput Screening. J. Am. Soc. Mass Spectrom. 2022, 33 (11), 2070–2077. 10.1021/jasms.2c00178.

(24) Radosevich, A. J.; Pu, F.; Chang-Yen, D.; Sawicki, J. W.; Talaty, N. N.; Elsen, N. L.; Williams, J. D.; Pan, J. Y. Ultra-High-Throughput Ambient MS: Direct Analysis at 22 Samples per Second by Infrared Matrix-Assisted Laser Desorption Electrospray Ionization Mass Spectrometry. Anal. Chem. 2022, 94 (12), 4913–4918. 10.1021/acs.analchem.1c04605.

(25) Pu, F.; Radosevich, A. J.; Bruckner, B. G.; Fontaine, D. A.; Panchal, S. C.; Williams, J. D.; Gopalakrishnan, S. M.; Elsen, N. L. New Platform for Label-Free, Proximal Cellular Pharmacodynamic Assays: Identification of Glutaminase Inhibitors Using Infrared Matrix-Assisted Laser Desorption Electrospray Ionization Mass Spectrometry. ACS Chem. Biol. 2023, 18 (4), 942–948. 10.1021/acschembio.3c00087.

(26) McInnes, L.; Healy, J.; Melville, J. UMAP: Uniform Manifold Approximation and Projection for Dimension Reduction. 2018. 10.48550/ARXIV.1802.03426.

(27) Moore, R.; Molyneux, C.; Sinclair, I.; Holdgate, G. A.; Walsh, J. Utilising Acoustic Mist Ionisation Mass Spectrometry to Identify Redox Cycling Compounds in High Throughput Screening Outputs. SLAS Discovery 2022, 27 (6), 369–374. 10.1016/j.slasd.2022.06.002.

(28) Xu, X.; Meng, Y.; Li, L.; Xu, P.; Wang, J.; Li, Z.; Bian, J. Overview of the Development of Glutaminase Inhibitors: Achievements and Future Directions. J. Med. Chem. 2019, 62 (3), 1096–1115. 10.1021/acs.jmedchem.8b00961.

(29) Gross, M. I.; Demo, S. D.; Dennison, J. B.; Chen, L.; Chernov-Rogan, T.; Goyal, B.; Janes, J. R.; Laidig, G. J.; Lewis, E. R.; Li, J.; MacKinnon, A. L.; Parlati, F.; Rodriguez, M. L. M.; Shwonek, P. J.; Sjogren, E. B.; Stanton, T. F.; Wang, T.; Yang, J.; Zhao, F.; Bennett, M. K. Antitumor Activity of the Glutaminase Inhibitor CB-839 in Triple-Negative Breast Cancer. Molecular Cancer Therapeutics 2014, 13 (4), 890–901. 10.1158/1535-7163.MCT-13-0870.

