## Supplemental for "Metabolite Fingerprinting for Phenotypic Screening by Infrared Matrix-Assisted Laser Desorption Electrospray Ionization Mass Spectrometry"

**Submitted to**: *bioRxiv*

**Submitted**: TBD

**Supplemental Manuscript:** 7 Supplemental Figures

**Keywords:** IR-MALDESI, phenotypic screening, drug discovery, cell-based assay, glutaminase, mass spectrometry

***Author for Correspondence**

Nathaniel L. Elsen, Ph.D.

Discovery Research

AbbVie Inc.

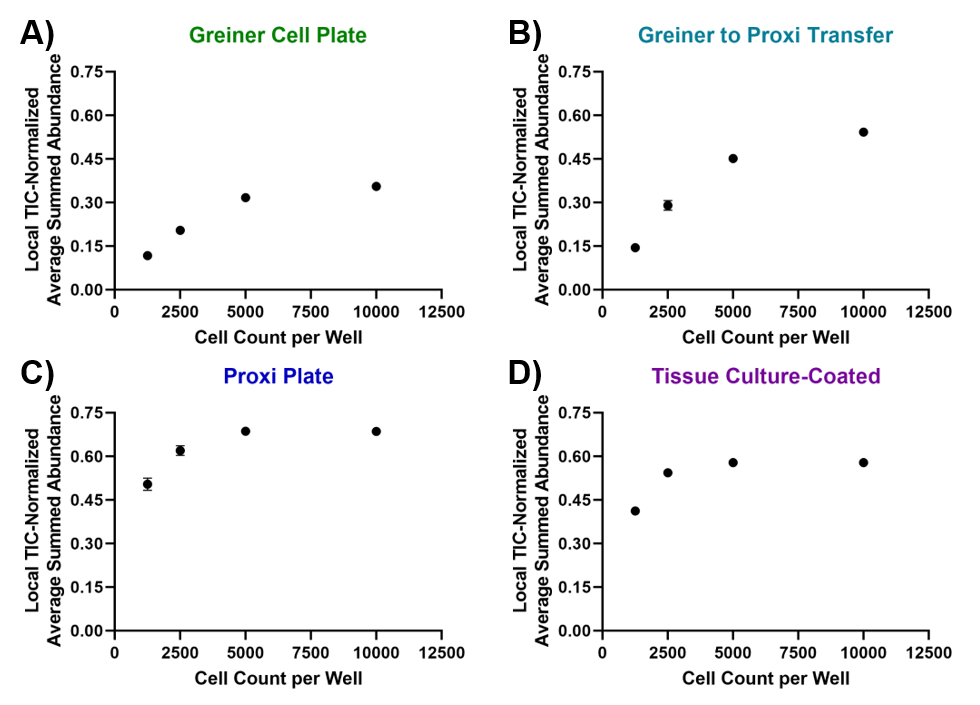

**Figure S1.** Summed metabolite abundances versus cell count in four conditions: **A)** plating on Greiner clear-bottom plate, **B)** plating on Greiner clear-bottom plate and transferring the cell lysate to a Proxi plate prior to MALDESI, **C)** plating directly on Proxi plate and **D)** plating directly on tissue-culture-coated Proxi plate. Local TIC bounds were *m/z* 100-200, 200-300, and 300-400 for normalization.

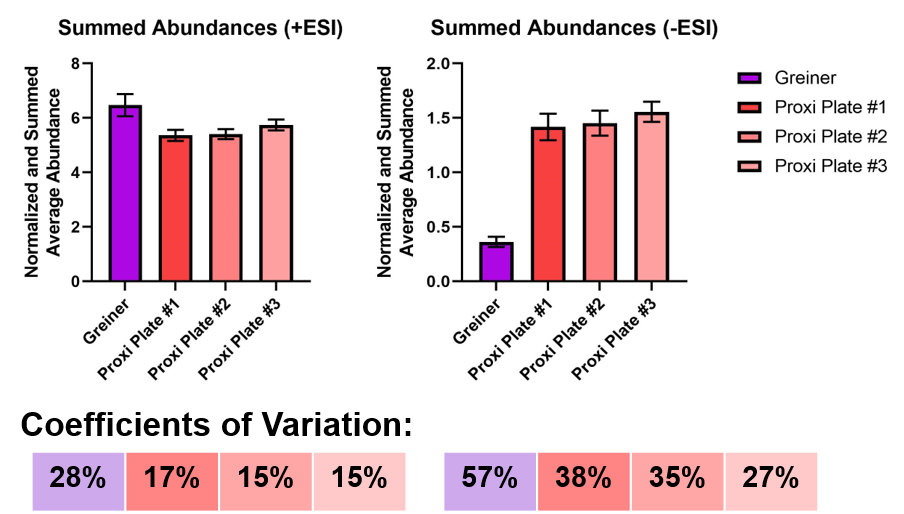

**Figure S2**. Summed average normalized abundances and % CV. Results obtained from analyzing lysate directly from the Greiner cell plate are in purple. Three aliquots were taken from a different Greiner plate and dispensed into Proxi plates in triplicate. Positive mode (+ESI) Proxi plate data was collected when the stage height was lower than optimal. Negative mode Proxi plate data is at the optimal stage height. Error bars correspond to the 95% confidence interval.

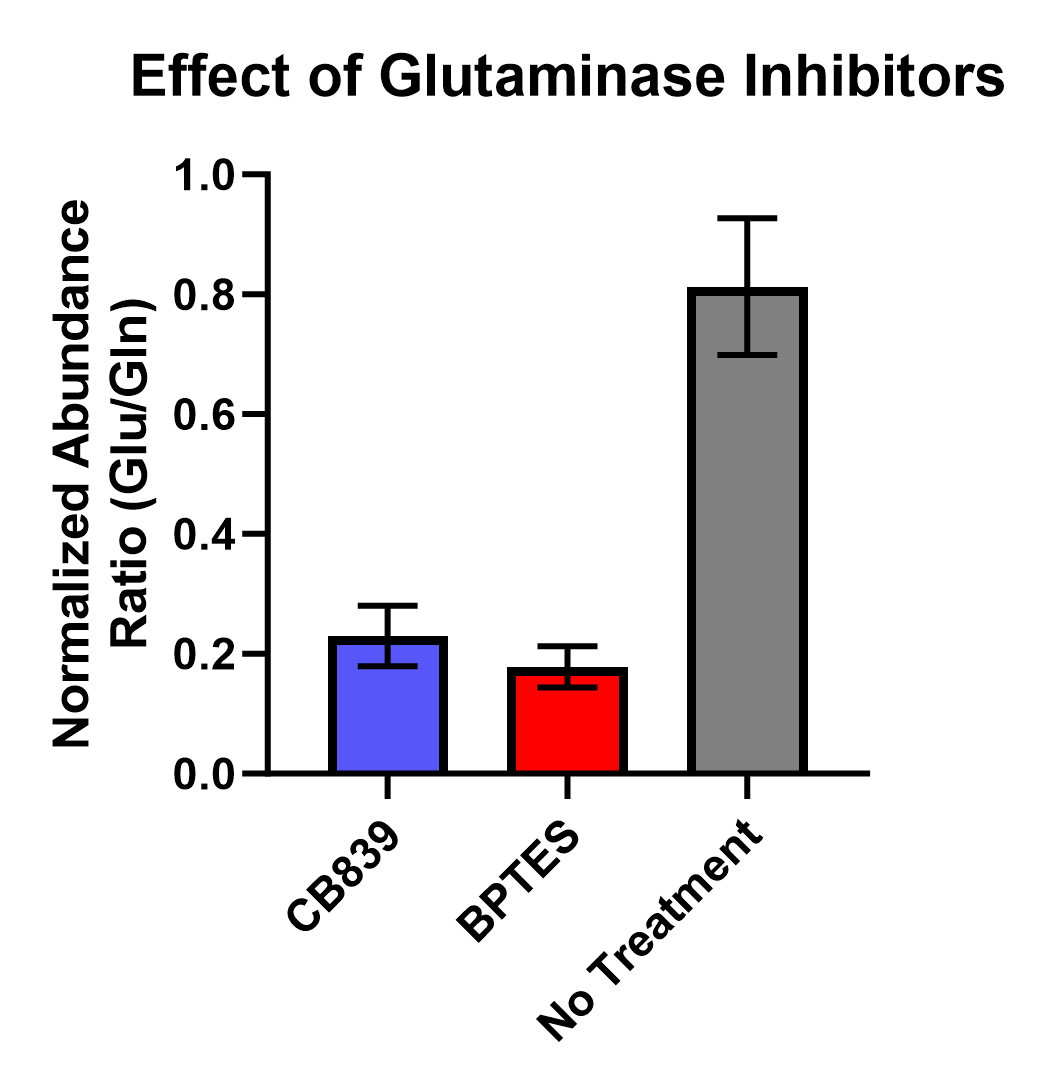

**Figure S3**. Average ratio of ^13^C_3_-caffeine-normalized abundances of glutamate (*m/z*: 148.0608, [M+H]^+^) to glutamine (*m/z*: 147.0764, [M+H]^+^) for CB839, BPTES, and No Treatment at 10 µM for four hours. Error bars correspond to the standard deviation.

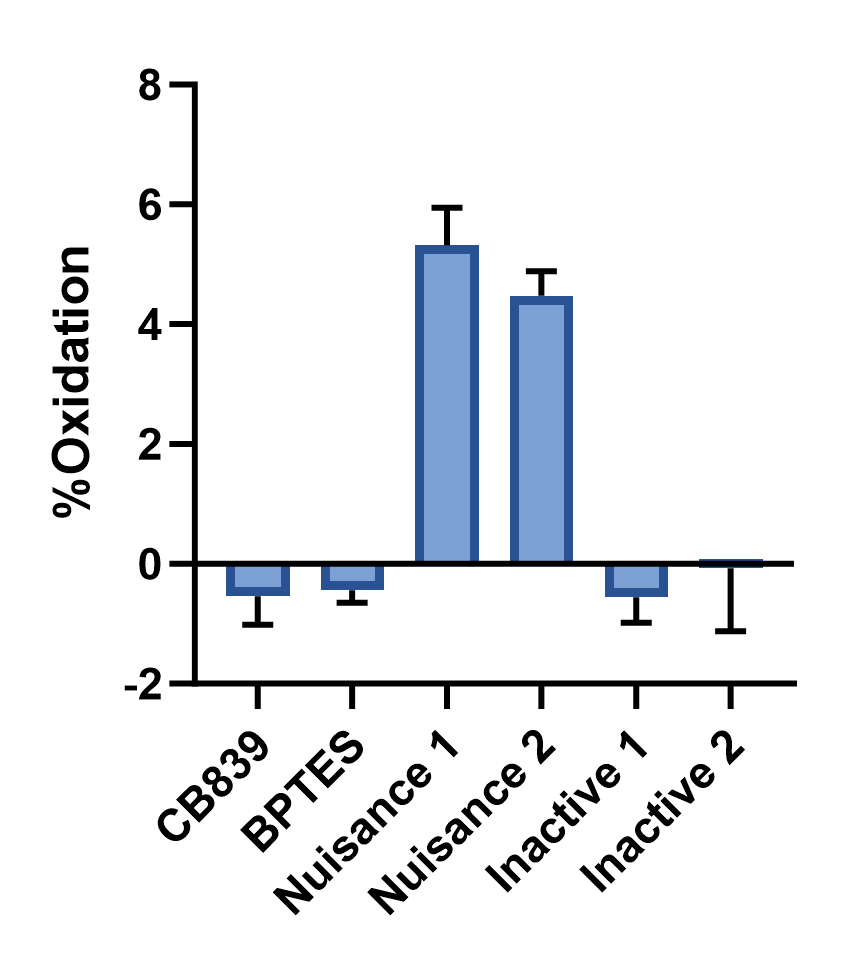

**Figure S4**. TCEP assay indicates the nuisance compounds have moderate level of redox activity. Error bars correspond to the standard deviation.

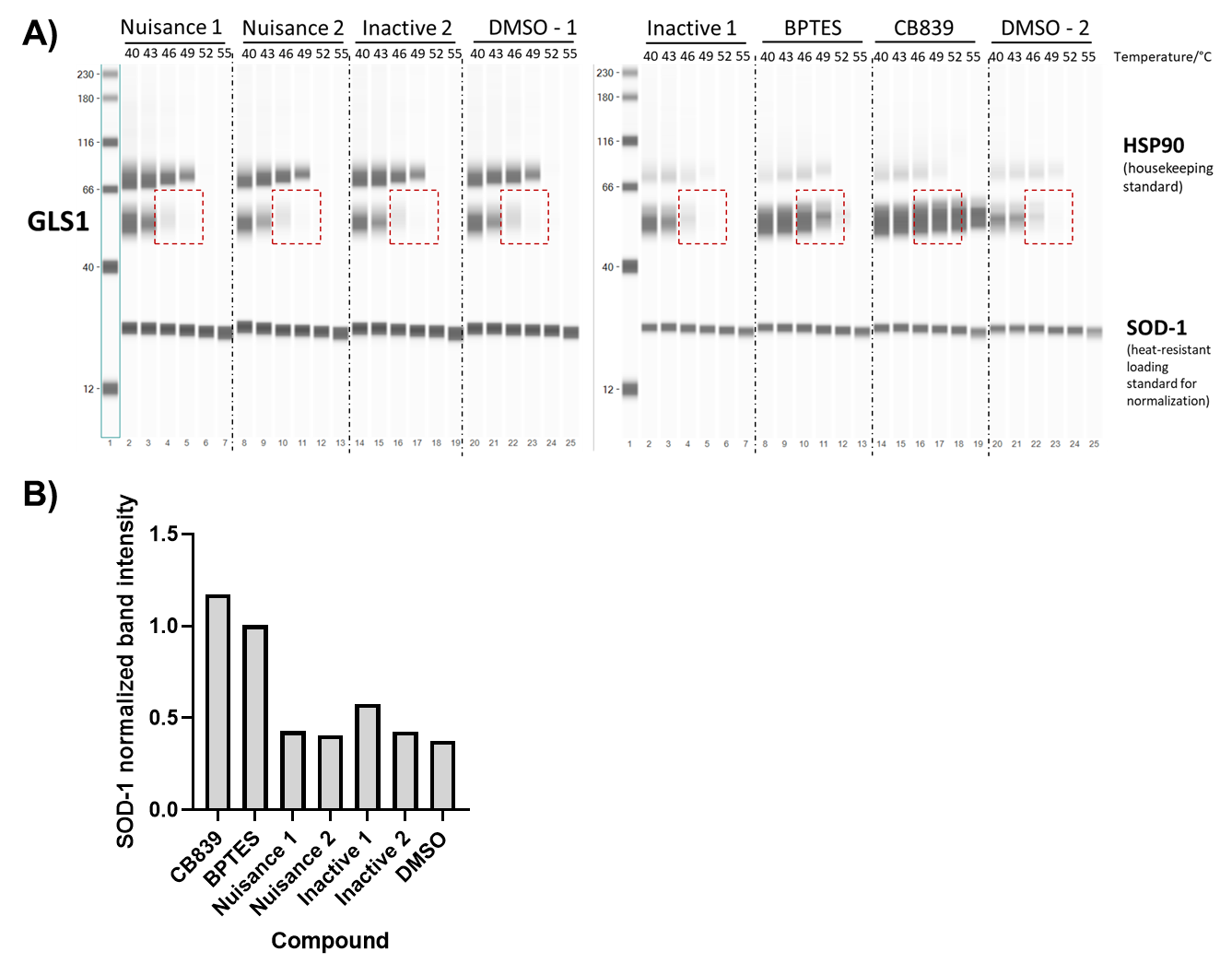

**Figure S5**. Cellular thermal shift assay (CETSA) indicates the two confirmed glutaminase inhibitors, CB839 and BPTES, showed a marked stabilization of GLS1 at 46 degrees while the other four compounds showed no stabilization. **A)** Raw western blot results. **B)** SOD-1 normalized band intensity at 46 °C.

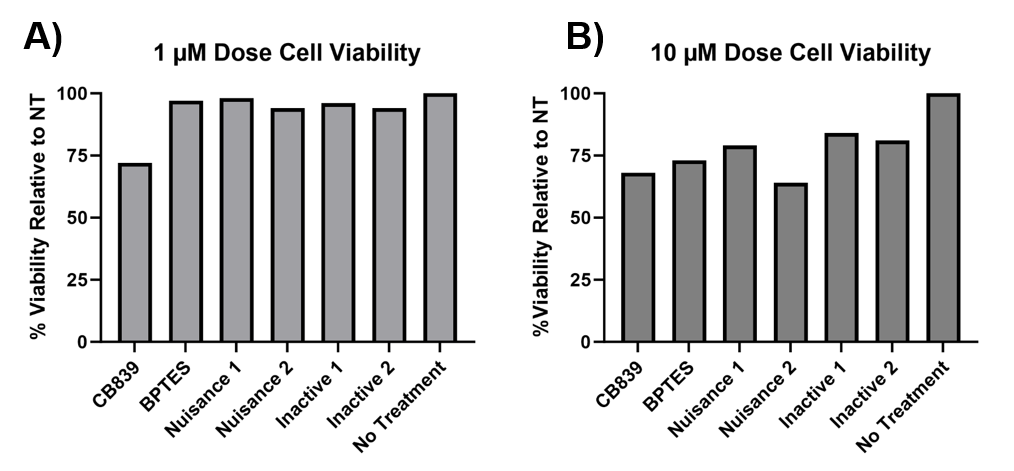

**Figure S6.** CellTiter-Glo summarized results for **A)** 24-hour incubation at 1 µM and **B)** 24-hour incubation at 10 µM. The raw luminescence was corrected for background and then scaled relative to the No Treatment control.

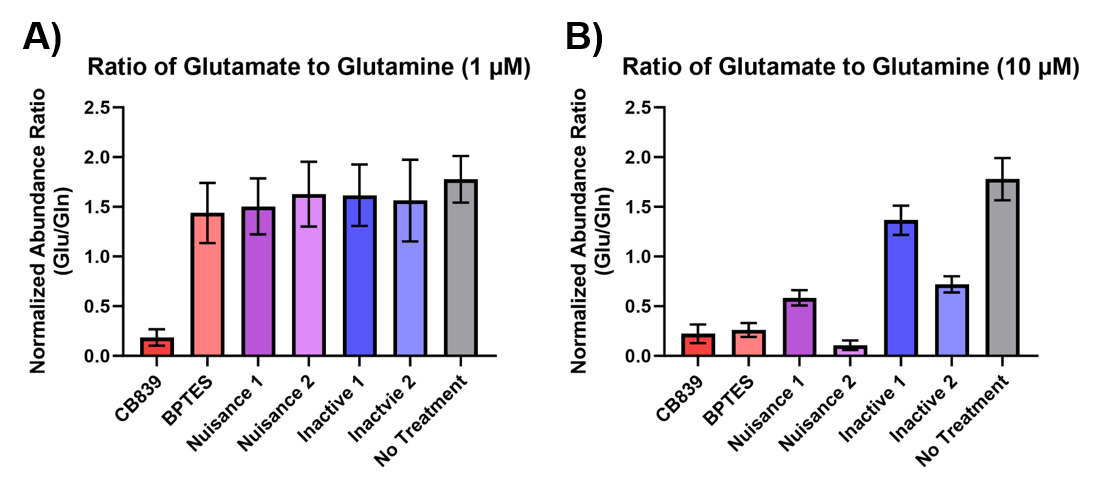

**Figure S7.** Average ratio of ^13^C_3_-caffeine-normalized abundances of glutamate (*m/z*: 148.0608, [M+H]^+^) to glutamine (*m/z*: 147.0764, [M+H]^+^) for all treatments at 10 µM for 24 hours. Error bars correspond to the standard deviation.
